# Diffusional and chemical control in the tyrosine kinase network of platelet CLEC-2 signalling

**DOI:** 10.1101/529859

**Authors:** Alexey A. Martyanov, Feodor A. Balabin, Joanne L. Dunster, Mikhail A. Panteleev, Jonathan M. Gibbins, Anastasia N. Sveshnikova

## Abstract

C-type lectin-like receptor 2 (CLEC-2) is platelet membrane glycoprotein implicated in maintenance of blood vessel integrity and development of lymphatics. Organization and regulation of tyrosine kinase signalling network associated with CLEC-2 is poorly understood. Here we aimed to investigate CLEC-2 signal transduction using computational systems biology methods in combination with experimental approaches.

We developed a 3D/stochastic multicompartmental computational model of CLEC-2 signalling, which was built around membrane-associated signalosome formation.

The model predicted that both Syk and Src-family kinases phosphorylate CLEC-2, and that CLEC-2 ligation induces cytosolic calcium spiking. This was experimentally confirmed using flow cytometry and TIRF microscopy, respectively. Sensitivity analysis suggested that the CLEC-2 translocation to the signalosome region is one of the rate-limiting steps in the signal transduction process. In agreement with this prediction, CLEC-2-induced platelet activation was strongly temperature-dependent (unlike that mediated by G-protein coupled receptors) and was delayed by lipid raft disruption.

Our results suggest a revised picture of the CLEC-2 signal transduction network functioning that emphasizes the crucial role of lipid raft structural rearrangement followed by tyrosine kinase feedback interplay.

## 1. Introduction

Prevention of the blood loss upon vessel wall disruption is the main task of platelets, the non-nucleated cellular fragments produced from megakaryocytes in the bone marrow (Versteeg *et al*, 2013). Platelets circulate in the cardiovascular system for approximately seven days, until they get eliminated in the spleen or liver (van der Meijden & Heemskerk, 2018). Alongside their primary role in hemostasis, platelets were demonstrated to be involved in angiogenesis (Repsold *et al*, 2017), tissue remodelling (Nurden, 2007; Gawaz & Vogel, 2013), and leukocyte recruitment under inflammatory conditions (Ed Rainger *et al*, 2015; Hitchcock *et al*, 2015). Platelets respond gradually to various activators, which occur during vessel wall injury (Versteeg *et al*, 2013). Thus, platelet response to extracellular matrix protein collagen includes several parts mediated by calcium signalling, namely shape change, granule release and, in some cases, cell death (Watson *et al*, 2010). In contrast, platelet response to ADP released from dying cells is brief and includes only shape change and integrin activation (Watson *et al*, 2010). To govern different responses to various activators, a platelet utilises its versatile signalling network capable of interpreting all potential extrinsic stimuli (Kauskot & Hoylaerts, 2012).

There are two main types of signalling pathways in blood platelets, associated with G-proteins and with tyrosine-kinase signalling (Stalker *et al*, 2012). Platelet G-protein coupled receptors (GPCR)/govern responses to ADP (P2Y_1_, P2Y_12_ receptors), thromboxane A_2_ produced by platelets upon stimulation (TP) activated during blood plasma coagulation thrombin (PAR1, PAR4), epinephrine (α2A) and produced by healthy endothelium prostacyclin (IP) (Gurbel *et al*, 2015; FitzGerald, 1991). Lesser types of platelet receptors induce a tyrosine-kinase network of signalling. There are receptors for collagen (GPVI) (Gibbins *et al*, 1997; Watson *et al*, 2010) and less known receptors for IgG (FcγRIIa) (Stalker *et al*, 2012). The most recently identified platelet receptor for lymphatic endothelium protein podoplanin CLEC-2 (C-type lectin-like receptor II-type (CLEC-2)) also transduces signals via tyrosine kinases (Watson *et al*, 2010).

CLEC-2 is a transmembrane signalling protein that induces platelet activation in response to a number of agonists: human cell plasma-membrane glycoprotein podoplanin (Suzuki-Inoue *et al*, 2007; Christou *et al*, 2008), snake venom protein rhodocytin (Huang *et al*, 1995; Shin & Morita, 1998), and brown seaweed extract fucoidan (Manne *et al*, 2013). The interaction of CLEC-2 with podoplanin, which is exposed on the surface of lymphatic endothelial cells (LECs), is crucial for the separation of blood and lymphatic systems during embryogenesis (Bertozzi *et al*, 2010; Suzuki-Inoue *et al*, 2010; Hughes *et al*, 2015) and for the prevention of blood-lymph mixing in high endothelial venules and lymph nodes in adult organisms (Herzog *et al*, 2013). Platelet CLEC-2 also contributes to the maintenance of blood vessel integrity during inflammatory conditions (Boulaftali *et al*, 2013; Hughes *et al*, 2010a; Bender *et al*, 2013; Gros *et al*, 2015; Hughes *et al*, 2015) and has a role in thrombus stabilization under flow conditions (May *et al*, 2009; Inoue *et al*, 2015; Hughes *et al*, 2010). Participation of platelet CLEC-2 has been demonstrated for a set of pathophysiological processes: promotion of tumor metastasis (Kato *et al*, 2008; Shirai *et al*, 2017), liver thrombosis after *Salmonella* Infection (Hitchcock *et al*, 2015), purpura and thrombocytopenia during Kazabach-Merritt syndrome in infants (O’Rafferty *et al*, 2015). CLEC-2 is also implicated in triggering deep vein thrombosis (Payne *et al*, 2017). Hence CLEC-2 was suggested to be a prospective therapeutic target (Chang *et al*, 2015; Payne *et al*, 2017; Hitchcock *et al*, 2015; O’Rafferty *et al*, 2015). Thus, systemic understanding of the CLEC-2 signalling is of essential importance.

CLEC-2 possesses a hemITAM signalling motif (which consists of single YxxL motif and is located in CLEC-2 cytoplasmic domain below negatively charged DED amino acid sequence) (Hughes *et al*, 2013). CLEC-2 molecules on the surface of platelets mostly exist as non-covalently bound dimers, which form larger clusters upon agonist ligation in the lipid rafts (Watson *et al*, 2009; Hughes *et al*, 2010b). Activated CLEC-2 transmits signals via the tyrosine kinase Syk and the Src family of tyrosine kinases (SFKs) (Pollitt *et al*, 2014). Syk and SFK control LAT-signalosome formation (Pollitt *et al*, 2010; Badolia *et al*, 2017; Manne *et al*, 2015b), which consists of PLCγ2, PI3K, SLP-76 and a set of other adaptor proteins. PLCγ2 activation leads to IP_3_ production and downstream calcium signalling. Pollitt et al. suggested that the primary platelet response to CLEC-2 ligation is weak and that CLEC-2 mediated platelet activation is largely due to the actions of secondary mediators of platelet activation: ADP and TxA2 (Pollitt *et al*, 2010).

The main feature observed upon platelet activation via CLEC-2 by either podoplanin, rodocytin or fucoidan was its 1-2 minute lag-time (Pollitt *et al*, 2010; Manne *et al*, 2013). The prolonged platelet CLEC-2 induced response may be a consequence of ADP-containing granule secretion and TxA2 synthesis and/or that of receptor clustering, as was proposed by Pollitt et al. (Pollitt *et al*, 2010). The platelet responses to CLEC-2 are also highly dependent on actin polymerization and cholesterol presence in the plasma membrane (Pollitt *et al*, 2010; Inoue *et al*, 1999). However, Badolia et al. have recently reported that CLEC-2 induced signalling is still detectable after abrogation of secondary activation (Badolia *et al*, 2017).

Here we aimed to understand the primary events of platelet activation upon stimulation through CLEC-2 utilizing mathematical modelling and experimental analysis. The proposed computational model describes platelet activation from ligand binding to CLEC-2 to calcium ions release from dense tubular system. Alongside the CLEC-2 model, the GPVI model has also been developed by modification of the initial stages of CLEC-2 induced platelet activation. This proves that differences between platelet CLEC-2 and GPVI models are in the initial stages of activation. The CLEC-2 model predicts that platelet activation via CLEC-2 is limited by the CLEC-2 ability to move in the plasma membrane. This was confirmed experimentally by an essential dependence of CLEC-2-induced platelet activation on temperature. CLEC-2 induced calcium spiking was predicted by the model and tested experimentally by means of TIRF-microscopy. Our data enable us to propose a novel view of the role of the features of the plasma membrane in signal transduction upon platelet activation via CLEC-2.

## 2. Results

### 2.1. CLEC-2 and GPVI model construction and validation

Our model scheme is given in Fig. 1 (CLEC-2) and Fig. S1 (GPVI). Both platelet CLEC-2 and GPVI receptors transduce signals via Immmunoreceptor tyrosine-based activation motifs: hemITAM and ITAM, respectively (Watson *et al*, 2010). A hemITAM contains single YxxL motif and an acidic amino-acid region DED above (Hughes *et al*, 2013), while ITAM consists of two YxxL motifs, separated by 6-12 amino-acids (Clemetson *et al*, 1999). An ITAM is located in the cytoplasmic region of the FcR γ-chain that is non-covalently linked to GPVI, CLEC-2 possess hemITAM directly in its cytoplasmic domain (Watson *et al*, 2010). After platelet activation via CLEC-2 or GPVI, a tyrosine residue in the hemITAM/ITAM sequence becomes rapidly phosphorylated by one of the tyrosine kinases, either Syk or representatives of the SFKs that are Lyn, Fyn, Src (Manne *et al*, 2015a; Hughes *et al*, 2010b, 2015). SFKs are initially presented in an autoinhibited state and can be half-activated by CD148 or PTP1B phosphatase (Senis *et al*, 2009; Mori *et al*, 2012). It has been reported that DUSP3 also participates in this initial activation of SFKs (Musumeci *et al*, 2015) which can localize to the plasma membrane via palmitoylation or association with proline-rich amino-acid sequences (GPVI cytoplasmic domain contains poly-Proline region) by its SH3 domain (Moroco *et al*, 2014; Bradshaw, 2010). Further SFK activation requires binding of its SH2 domain to the phosphorylated tyrosine residues in the hemITAM/ITAM region (Bradshaw, 2010). Unlike SFKs, Syk does not exhibit gradual activation. Syk activation can be performed via two pathways: binding of both Syk SH2 domains to phosphorylated tyrosine residues in hemITAM/ITAM or phosphorylation of the tyrosine in the linker region between Syk kinase domain and SH2 domains. The linker region has been shown to be phosphorylated by active SFK or Syk (Bradshaw, 2010; Tsang *et al*, 2008; Hughes *et al*, 2015).

**Figure 1.**
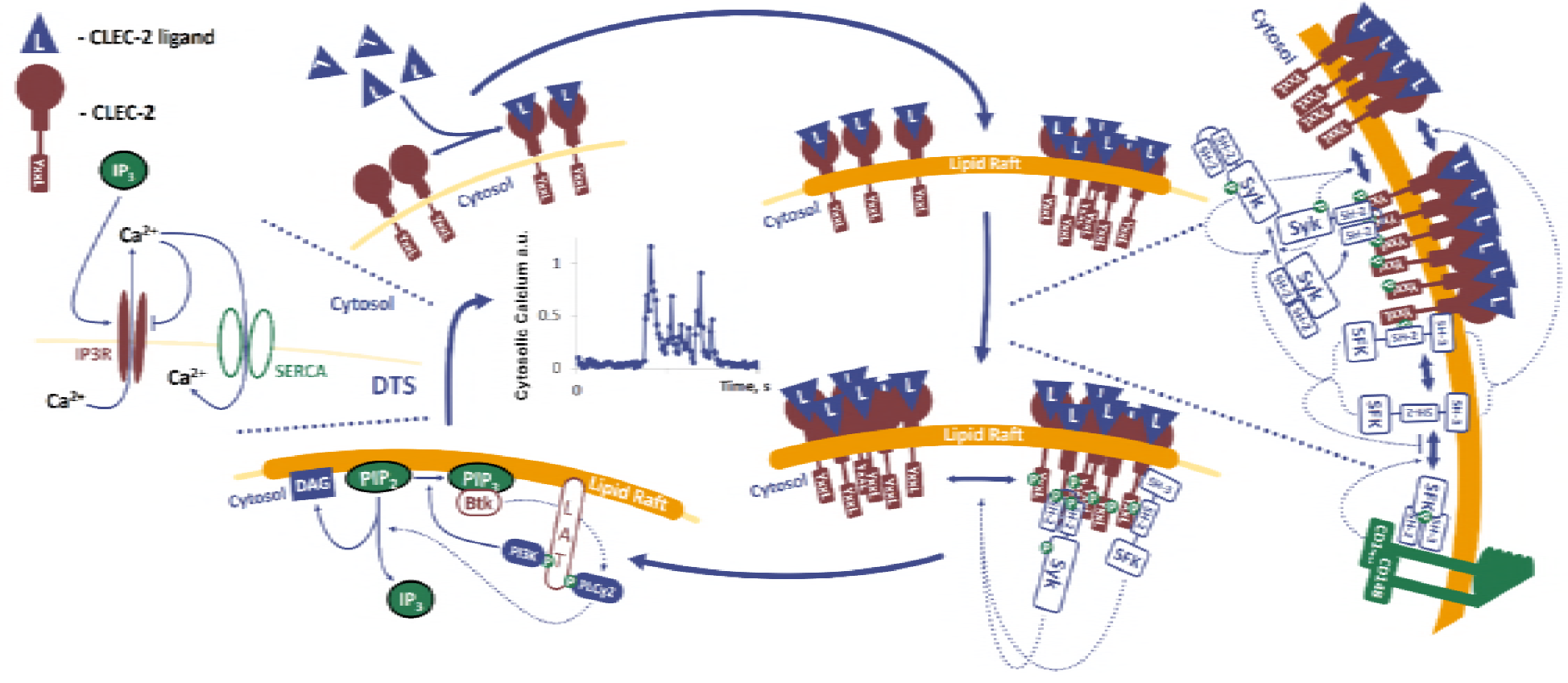
Scheme of CLEC-2 induced signalling in a modular fashion. Initially active CD148 produces half-active SFK. Active forms of SFK can phosphorylate Syk on Y346 converting the enzyme to an active state. Following initial activation, Syk can also auto-phosphorylate. Upon ligation of CLEC-2 and fucoidan, translocation of the activated CLEC-2 to lipid rafts occurs. In lipid rafts, CLEC-2 molecules form clusters. Active SFK and Syk phosphorylate hemITAM in the CLEC-2 cytoplasmic domains. Inactive Syk or half-active SFK bind two (one for SFK) phospho-hemITAMs with their SH-2 domains and become active. Active Syk phosphorylates the adaptor protein LAT. PLCɣ2 and PI3K bind P-LAT. This activates PI3K, which phosphorylates PIP_2_ and produces PIP_3_, which becomes a docking site for Btk. Btk becomes activated and can in-turn activates PLCɣ2. Active PLCɣ2 hydrolyses PIP_2_ to IP_3_ and DAG. IP_3_ binds to IP3R on the surface of the dense tubular system (DTS). This activates IP3R. Through active IP3R free Ca^2+^ ions pass to the cytosol. Ca^2+^ can inhibit IP3R as well as return to the DTS via SERCA.

Once bound to ligands, GPVI and CLEC-2 receptors are translocated to the lipid rafts - plasma membrane microdomains rich in cholesterol (Pollitt *et al*, 2010; Watson *et al*, 2010). The exact mechanism of the CLEC-2 and GPVI receptor clustering is yet unknown. In the lipid rafts, receptor molecules are brought to the close proximity with a variety of signalling proteins, among which are SFK, Syk, LAT, PI3K, PLCγ2 (Stalker *et al*, 2012; Hughes *et al*, 2010b; Pollitt *et al*, 2014). Thus, lipid rafts govern intracellular signalling upon receptor tyrosine-kinase stimulation in platelets.

The main differences between CLEC-2 and GPVI signalling are in the primary activation events described above. As soon as Syk tyrosine kinase is activated, GPVI and CLEC-2 signalling pathways are assumed to coincide (Stalker *et al*, 2012; Watson *et al*, 2010).

We developed 3D reaction-diffusion and 0D stochastic mechanism-driven computer models that capture the regulation of CLEC-2 and GPVI induced platelet activation. The systems of equations that form the computational models were constructed from current biological knowledge of the biochemical reactions utilising assumptions of either mass action or Henry-Michaelis-Menten. Parameter values were taken from experimental reports on the respective human enzymes while the numbers of proteins per platelet were taken from published proteomics (Burkhart *et al*, 2012). A detailed description of the computational model and its versions can be found in the supplemental material, where, due to the complexity of CLEC-2 signalling, we divide the full model into biologically relevant modules (see also Fig. 1). The detailed model description including model reactions and parameters can be found in Tables S1-S17. The full 3D reaction-diffusion model is a set of 19 partial differential equations and 62 parameters, with 50 parameters obtained from the literature. The full stochastic model describes the behavior of 53 species in 87 reactions and contains 53 independent parameters, with 21 parameter values obtained from the literature. The geometry of the spatial model consists of three compartments: extracellular volume, plasma membrane and cytosol. To decrease the computational time, the spatial model was converted into a homogenous (0D) model in the following manner. The diffusion was considered as a translocation of species from one spatial location to another and thus the reaction model volume was divided into two sub-volumes between which the species translocate. The similarity of the outcome of these two representations was demonstrated computationally on Figure S2. The geometry of the stochastic model consists of six compartments: extracellular volume, plasma membrane adjacent volume, lipid raft adjacent volume, cytosol, DTS adjacent volume and DTS. Geometric regions are adjusted to describe cytosol-membrane to volume ratio observed in platelets (Eckly *et al*, 2016), details are given in Table S2, S10. Initial concentrations are given in Tables S2-S4, S6, S7, S11, S13. Due to the complexity that has been unveiled in CLEC-2 induced signalling in platelets the scheme of biochemical processes was considered in a modular fashion (Fig. 5, Tables S5, S8, S12, S14). A detailed description of parameter estimation process is given in supplementary materials.

CLEC-2 model incorporates both known exogenous CLEC-2 ligands: fucoidan and rhodocytin. Fucoidan can bind only single CLEC-2 molecules, while rhodocytin is capable of binding a CLEC-2 dimer (Ustyuzhanina *et al*, 2016; Watson *et al*, 2008). However, while fucoidan is a polysaccharide, rhodocytin is a tetramer (Watson *et al*, 2008) and thus, diffusion speed of the CLEC-2-fucoidan complexes is significantly higher than diffusion speed of the CLEC-2-rhodocytin complexes.

Receptor clustering in the 3D model is simulated by a region in the membrane, lipid raft, where receptor diffusion constant is significantly decreased. For the stochastic model, receptor clustering was simulated by equations based on the law of mass action. Details are given in the supplement.

As soon as activated CLEC-2 molecules are translocated to the lipid raft, the hemITAM in the cytoplasmic domain of an activated CLEC-2 is phosphorylated by Syk (captured in “Syk-only” model) or Syk and SFK (“Syk-SFK” model). This reaction only occurs only in the signalling region (see details in Tables S1, S5, S12). Initially 8-10 active Syk (out of a total of 4900 molecules) and 300 of half-active SFK (out of a total of 36800 molecules) are present in the system. This is due to the presence of CD148, which dephosphorylates SFK and removes it from the autoinhibited state (Mori *et al*, 2012; Rollin *et al*, 2012). These half-active SFK can, in turn, activate small amounts of Syk by phosphorylating them at the linker region and turning them into an active state (Hughes *et al*, 2015).

As soon as CLEC-2 molecules are phosphorylated, non-active Syk kinases bind to them via SH2 domains and become phosphorylated and activated. This leads to the LAT phosphorylation by Syk and signalosome assembly: LAT associates with PI3K and PLCγ2. PI3K becomes active and produces PIP_3_ from PIP_2_. PIP_3_ is rapidly bound by PH-domain of the Btk kinase, which becomes active, and in turn activates PLCγ2. Finally, PLCγ2 hydrolyses PIP_2_ and produces IP_3_. IP_3_ production induces Ca^2+^ spiking. The model of Ca^2+^ release is described in our previous work (Sveshnikova *et al*, 2016; Balabin & Sveshnikova, 2016), and all reaction are incorporated in the model as depicted in Fig. 1.

We compared predictions of “Syk-SFK” and “Syk-only” versions of the model with published experimental data on protein tyrosine phosphorylation after activation of human platelets by fucoidan (Manne *et al*, 2013) or after activation of murine (Hughes *et al*, 2015) or human (Musumeci *et al*, 2015) platelets by rhodocytin (Fig. 2). The parameters of the model were not adjusted to murine platelet proteomic data and the achieved description of the experimental data was unexpected. The “Syk-only” model failed to describe both sets of available experimental data, while the “Syk-SFK” model was in good agreement with all data (Fig. S3).

**Figure 2:**
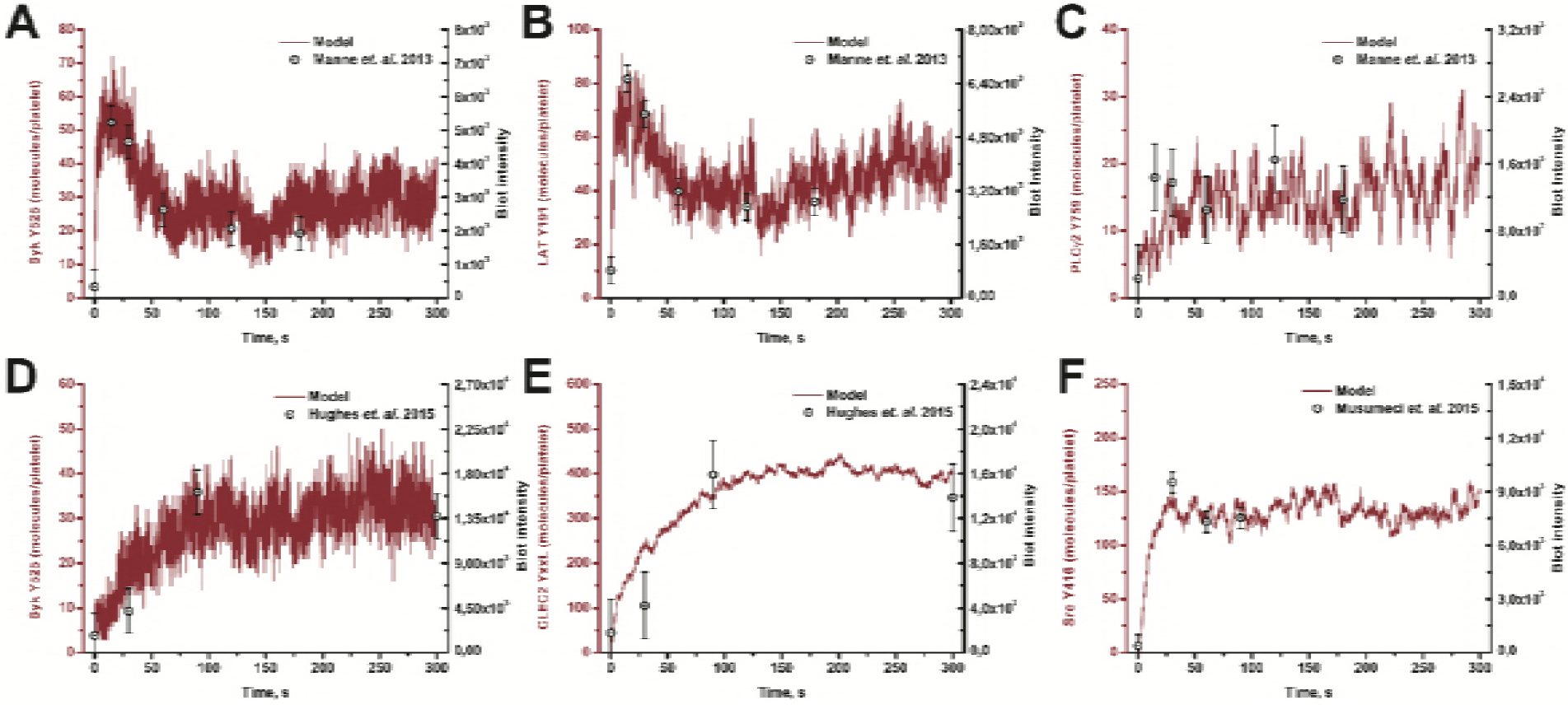
Computational model validation on published protein phosphorylation data. Stochastic model simulation (grey lines), with typical runs out of n=100 are given. Experimental data (black dots) were assessed using ImageJ software from the indicated published western-blotting data. (A-C) Time-dependent phosphorylation of participants of CLEC-2 signalling cascade after activation by 50 μg/ml of fucoidan. Experimental data are based on western blotting assays from (Manne *et al*, 2013), where human platelets were utilized. (D,E) Time-dependent phosphorylation of participants of CLEC-2 signalling cascade after activation by 300 nM of rhodocytin (Hughes *et al*, 2015). Experimental data in (F) were based on western blotting assays from (Musumeci *et al*, 2015). Murine platelets were utilized in these experiments.

For GPVI-associated signalling we modified the stochastic CLEC-2 model to incorporate the known differences between these signalling pathways. The GPVI model differs in the following way: the number of GPVI receptors were set to 5000 per platelet (Dunster *et al*, 2015) and the GPVI cytoplasmic tail was assumed to be phosphorylated only by SFK (utilising the same kinetic parameters as for CLEC-2). The rate constant of the ligated GPVI translocation to lipid raft region and GPVI dephosphorylation was adjusted to describe experimental data on Syk-Y525 phosphorylation following platelet stimulation with collagen related peptide (Dunster *et al*, 2015) and our own data describing cytosolic calcium dynamics following platelet stimulation with CRP in flow cytometry (Fig. S1).

### 2.2. Sensitivity analysis of the models: receptor diffusion and tyrosine-kinase activity have similar effect of platelet CLEC-2 induced activation

To find the critical controlling steps in CLEC-2 and GPVI induced platelet activation we performed a local sensitivity analysis (for deterministic simulation) of the constructed mathematical model as well as in the similar GPVI-induced platelet activation model (Fig. 3, Table S17). The activity of PLCγ2 after 20 s appeared to be most sensitive to changes in the number of CLEC-2 copies, activity (catalytic constants) of Syk and SFK kinases, LAT availability, and activities of PI3K and Btk. We performed a detailed investigation of the influence of these parameters on the model behavior (Fig. 3, S4-5). The parameters concerning phosphorylated CLEC-2 concentration in the signalling region were the most influential ones. Although molecular mechanisms of the assembly of the receptor-clusters remain unclear (Pollitt *et al*, 2010), in the model we assumed that the clusters of phosphorylated CLEC-2 are formed in the area close to LAT signalosome (LAT mostly exists in the lipid rafts and can be used as a specific marker of these regions (Pollitt *et al*, 2010)). This led to the theoretical prediction that the lag-time of platelet activation in response to CLEC-2 agonists should be dependent on the rate of translocation of CLEC-2 molecules (Fig. 3A, S4, S5) into the lipid rafts. Ligand affinity also affects the time to activation upon CLEC-2 stimulation (Fig. 3B), however, its sensitivity score is significantly lower than the sensitivity scores of other model parameters (Table S14).

**Figure 3:**
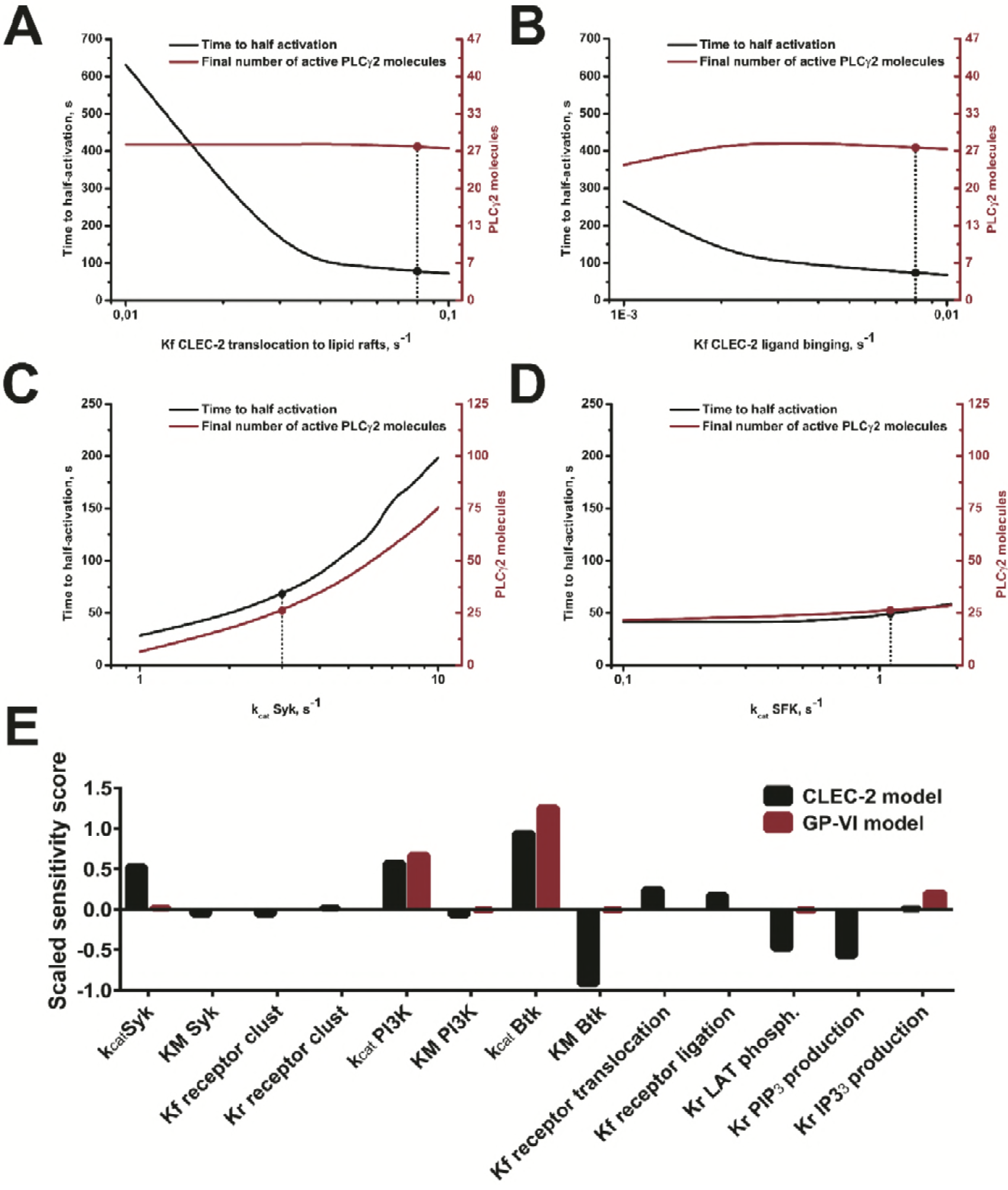
The rate-limiting steps in platelet tyrosine-kinase-mediated activation. Activity of PLCγ2 is used as a marker of platelet activation. Time to half-activation and maximum activity of PLCγ2, obtained after fitting with Hill-equation (see supplement), for variation of translocation rate (diffusion coefficient) of CLEC-2 molecules (A); ligand affinity (B), and activity rates (k_cat_) of Syk and SFK kinases (C, D). (E) Sensitivity analysis (utilizing the activity of PLCγ2 at 20s as a sensitivity score) of the deterministic models following 100 μg/ml of fucoidan (for CLEC-2 model) or 10 μg/ml of CRP (for GPVI model). Only the most sensitive parameters are shown here; the sensitivities to changes in all parameters are given in Table S14.

In order to investigate the influence of the size of the lipid raft on CLEC-2 induced signalling, spatial simulations were performed. While the general size of lipid rafts can be varied in the stochastic model, the size of individual rafts could not be assessed and therefore the model was modified to include spatial interactions (3D reaction-diffusion model), allowing assessment of the size of a single lipid raft on signalling (Fig. S6). Increasing the size of the pre-existing raft decreased lag-time before activation non-significantly for Syk-SFK model (Fig. S6 A), while for Syk, the model effect was more substantial (Fig. S6 B).

Kinetic parameters capturing the activity of the main signalling tyrosine kinases Syk, SFK and Btk had a similar impact on platelet activation when compared to the rate of receptor translocation (Fig. 3C,D). Further investigation (Fig. S7) (of the stochastic model) without a distinct signalling region resulted in a lack of platelet activation in response to CLEC-2 agonists. Thus, we conclude that, theoretically, membrane diffusion and CLEC-2 concentration in lipid rafts are the rate-limiting factors under normal conditions.

### 2.3. Experimental validation: CLEC-2 induced Ca^2+^ release is highly dependent on Syk and SFK kinases

An unresolved key question about the mechanisms of CLEC-2 induced platelet signalling is whether both Syk and SFK phosphorylate CLEC-2 cytoplasmic domain, or whether Syk is solely responsible. There are multiple reports that upon treatment of platelets with selective Syk inhibitors PRT060318 (Hughes *et al*, 2015) and R406 (Spalton *et al*, 2009) CLEC-2 phosphorylation is impaired. Moreover, Severin et al. 2011 demonstrated that platelet activation by CLEC-2 ligands is not affected after depletion of two of the three major SFK members (*fyn^−/−^*/*lyn^−/−^* or *lyn^−/−^*/*src^−/−^* or *fyn^−/−^*/*src^−/−^*) or depletion of CD148 phosphatase, which is one of the mediators of SFK activation (Severin *et al*, 2011). Based on these results, Severin et al. reported that CLEC-2 phosphorylation is mediated solely by Syk. However, authors did not perform experiments on platelets deficient in all of the SFK members, and these kinases might be interchangeable. The hypothesis that PTP1B may substitute for CD148 was not considered in (Mori *et al*, 2012). Double depletion of CD148 and PTP1B completely abrogated platelet responses to CLEC-2 agonists (Mori *et al*, 2012). These data together with the fact that broad SFK inhibitor PP2 disrupts CLEC-2 phosphorylation in a manner closely resembling PRT060318 (Hughes *et al*, 2015), show that the role of SFK in platelet CLEC-2 signalling is not obvious. Spalton et al. 2009 claimed that SFKs might perform initial phosphorylation of CLEC-2 while Syk enhances this process (Spalton *et al*, 2009). On the other hand, Hughes et al. 2015 proposed an opposite concept (Hughes *et al*, 2015).

To test if both Syk and SFK kinases are significant for platelet activation, hirudinated whole blood with Fura-RED loaded platelets was incubated with either Syk kinase inhibitor PRT060318, SFK kinase inhibitor PP2, or vehicle for control, activated with fucoidan for 0-5 min, diluted 40 times and directly analyzed by flow cytometry (Fig. 4A,B,C). While significant cytosolic calcium mobilization was observed in response to both 10 and 100 ug/ml fucoidan without inhibitors (Fig. 4B,C), any of the inhibitors significantly reduced this response. This is in line with the model prediction that both kinases are necessary for CLEC-2 induced platelet activation (Pollitt *et al*, 2010) (Fig. 4D,E).

**Figure 4.**
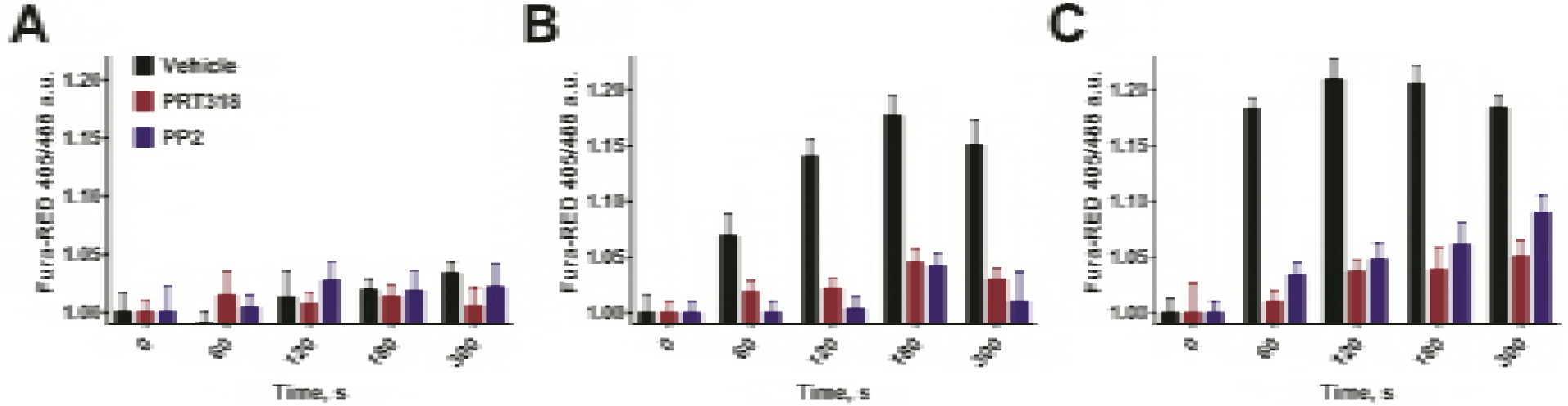
Flow cytometry assay of platelet cytosolic calcium mobilization in whole blood induced by fucoidan at 1 (A), 10 (B) or 100 (C) μg/ml in presence of Syk (PRT060318, 5 μM) or SFK (PP2, 20 μM) inhibitors. DMSO was used as vehicle control.

### 2.4. Experimental confirmation: temperature variation has a significant impact on platelet activation via CLEC-2

In order to assess the influence of the secondary mediators of platelet activation of CLEC-2 induced signalling, we pre-incubated platelet with P2Y12 receptor antagonist MRS2179. This had no effect on fucoidan-induced CLEC-2 activation, while activation by rhodocytin was disrupted (Fig. 5 B). Thus, in order to observe the signal upon CLEC-2 stimulation uninfluenced by secondary mediators, fucoidan was utilized in all further experiments. These data are in agreement with the fact that CLEC-2 can induce signalling independently from secondary mediators (Badolia *et al*, 2017). The influence of membrane saturation by cholesterol was studied using cholesterol binding agent mβCD (Mahammad & Parmryd, 2015). While high concentrations (5 mM) of mβCD were capable of complete abrogation of platelet response to fucoidan, relatively low concentrations (1 mM) delayed activation not affecting its degree (Fig. S8 B). Thus, 1 mM of mβCD was utilized for testing of the role of lipid raft size in all further experiments.

**Figure 5.**
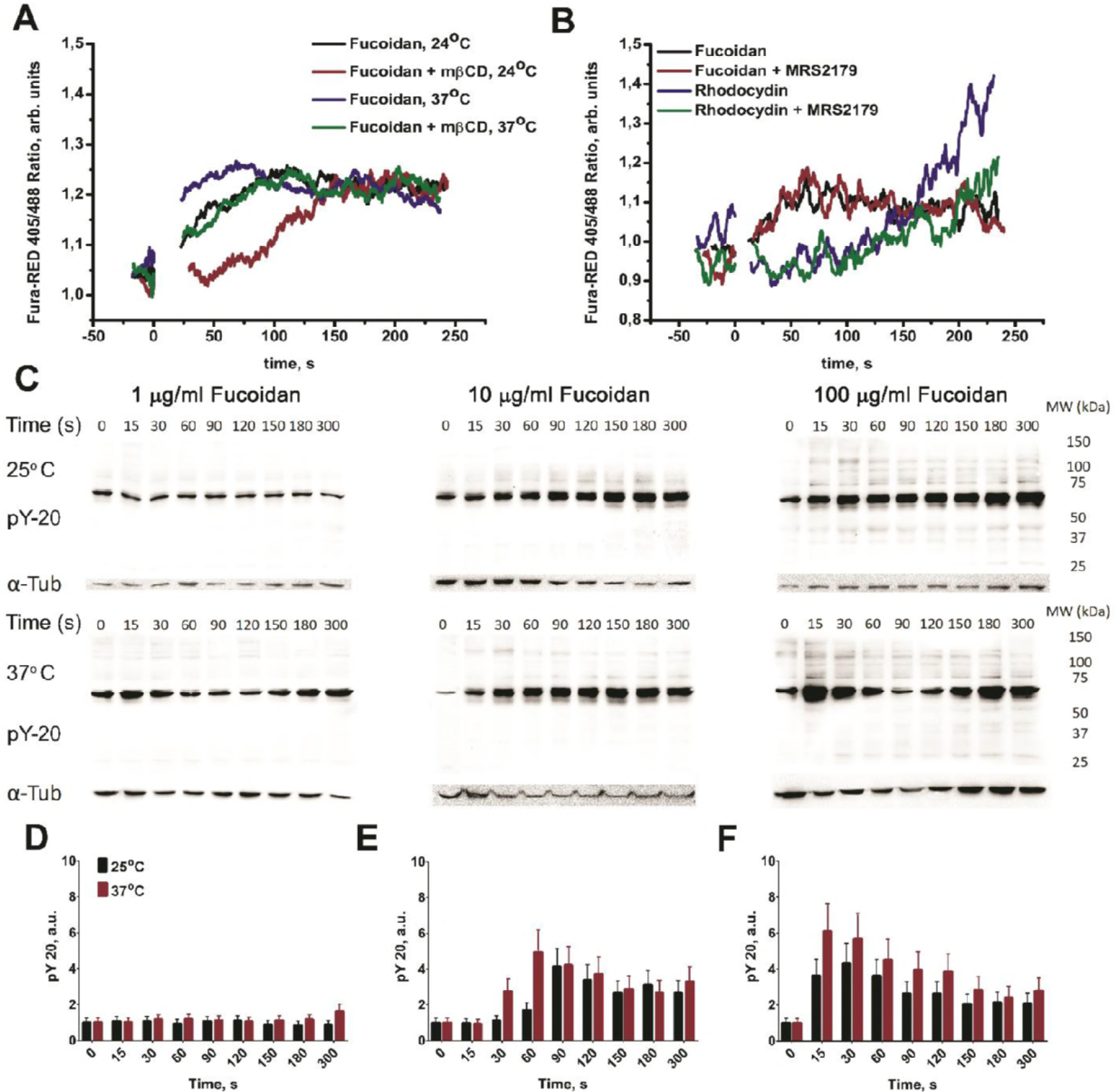
A – flow cytometry assay of CLEC-2 induced signalling in platelets after activation by 100 μg/ml of fucoidan. Activation at 25°C was significantly slower than at 37°C. Disruption of lipid rafts by mβCD delayed activation. Each curve represent data from at least three experiments. Ca^2+^ response is dependent on temperature conditions as well as on cholesterol saturation. B – Pre-incubation of platelets with MRS2179 had no effect on platelet response to 100 μg/ml fucoidan, while activation was disrupted upon stimulation by rhodocytin. C,D,E,F – Immunoblot assay of CLEC-2 induced signalling (C – raw data): platelets were activated by 1(D)-10(E)-100(F) μg/ml of fucoidan at 25°C or 37°C. Samples for analysis were taken 0-15-30-60-90-120-150-180-300 s after activation. Murine anti-human-phosphotyrosine primary anti-bodies (PY-20 clone) were used. Anti-tubulin primary antibodies were used as loading control after stripping. Typical experiment for one out of n = 3 different donors.

At both room (25°C) and body (37 °C) temperature, fucoidan increased cytosolic Ca^2+^ concentration (Fig. 5 A). However, at 25°C this increase occurred at 100 s, while at 37 °C it was significantly more rapid and occurred at 60 s. Pre-incubation of platelets with mβCD further delayed the increase in the concentration of cytosolic calcium both at 25 °C (240 s) and at 37 °C (105 s); the fact that degree of the response does not change suggests that lipid raft formation is not simply essential, but rather rate-liming in this phenomenon. The demonstrated influence of the temperature conditions on platelet activation by fucoidan supports the model prediction that the receptor translocation rate in the plasma membrane is one of governing parameters of the CLEC-2 induced platelet activation (Fig. 5 A). It should be noted that temperature variation had no effect on activation of platelets by low concentrations (2 μM) of ADP (Fig. S8 A).

To observe whether temperature influences all parts of the CLEC-2 signalling cascade, we performed immunoblotting analysis of tyrosine phosphorylation level in platelets activated by fucoidan at room and body temperature (Fig. 5 C-F). Washed human platelets at concentration 1.5 10^9^/ml in modified Tyrode’s buffer (no BSA and no Ca^2+^) were incubated with fucoidan for 0-5m at given temperature conditions. Samples were taken at 15-30 s time intervals and analyzed by immunoblotting with anti-phospho-tyrosine antibody PY20 as described in Methods section. The analysis show that while 1 μg/ml of fucoidan does not induce significant activation, both 10 and 100 μg/ml of fucoidan induced tyrosine phosphorylation with maximum after 90 s and 30 s respectively, and the increase in solution temperature significantly shortened the lag-times to 60 s and 15 s correspondingly.

### 2.5. Platelet CLEC-2-induced calcium spiking is predicted theoretically and demonstrated experimentally

We performed stochastic calculations to investigate the primary cytosolic calcium signalling in platelets stimulated with fucoidan in two versions of our model. The “Syk-only” model restricts CLEC-2 to be phosphorylated by Syk kinase; the “Syk-SFK” model, allows CLEC-2 to be phosphorylated by both Syk and SFK kinases (see Fig. 6 for a comparison of model outputs). In response to fucoidan, both models predict cytosolic calcium spiking (Fig. 6A,C) and similar increases in average cytosolic calcium concentration, of 60-100 nM (Fig. 6B,D). Upon rhodocytin stimulation only “Syk-SFK” model predicted significant cytosolic Ca^2+^ concentration change (Fig. 6F-I) at various receptor diffusion, clustering and dephosphorylation rates. Variation of the kinase activity did not restore spiking for “Syk-only” model. As can be seen from Fig. 6A,F, the “Syk-SFK” model predicts that CLEC-2 can induce Ca^2+^ spiking after activation by either fucoidan or rhodocytin. However, the “Syk-only” model predicted a delayed Ca^2+^ spiking upon fucoidan stimulation (Fig. 6C) and no cytosolic calcium spiking after stimulation by rhodocytin (Fig. 6H). Thus, according to our calculations, primary activation by both fucoidan and rhodocytin can induce cytosolic calcium mobilization in platelets. In addition, the data on Fig. 6 further support the thesis that both Syk and SFK may phosphorylate CLEC-2. Similar results were obtained after averaging results of 500 calculations (Fig. 6B,G).

**Figure 6.**
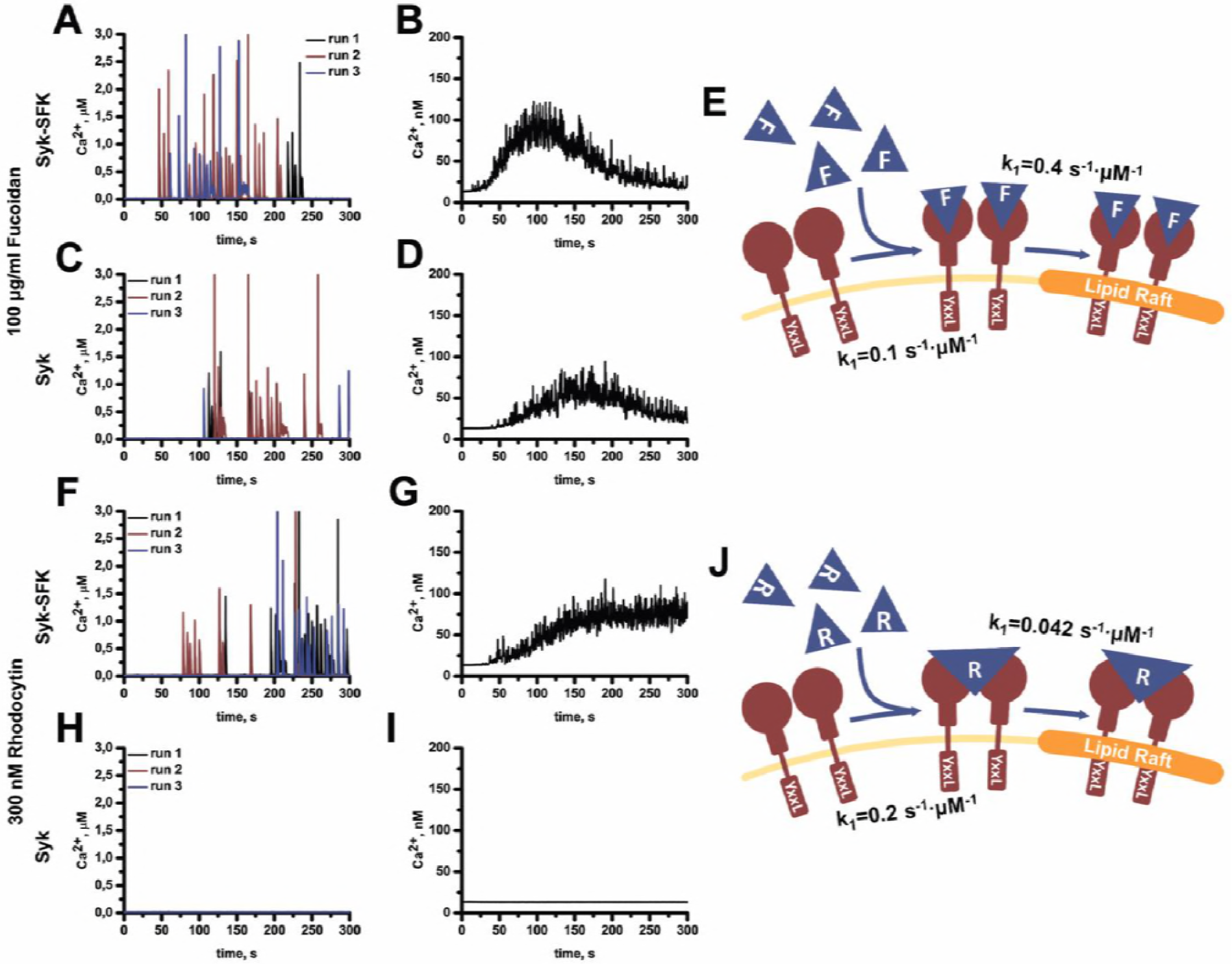
Stochastic simulation of platelet activation through CLEC-2. Stochastic simulation of cytosolic calcium levels in platelets stimulated with 100 μg/ml fucoidan (A-D) or 300 nM rhodocytin (F-I) for “Syk-SFK” (A,B,F,G) or “Syk-only” (C,D,H,I) models. For each condition three typical runs out of n = 100 are given (A,C,F,H). Calcium concentration was averaged upon 500 stochastic runs (B,D,G,I). Schematic representations of the reactions underlying CLEC-2 activation with Fucoidan (E) and Rhodocytin (J) are given.

To investigate the nature of the cytosolic calcium increase observed in flow cytometry total internal reflection fluorescent (TIRF) microscopy of immobilized single calcium-sensitive dye loaded platelets was performed. We utilized two experimental settings. In the first setting platelets were immobilized on VM64 (anti-CD31 clone (Mazurov *et al*, 1991)) antibody (Fig. 7A-D) and fucoidan solution in Tyrode’s buffer was washed over the surface. In the second setting, fucoidan was applied to hydrophobic cover glass and then a washed platelet suspension in Tyrode’s buffer was perfused over the glass (Fig. 7E-H). In both settings, platelets became attached to the surface and developed filopodia (Fig. 7 B, D, F, H). In both cases, platelets developed cytosolic calcium spiking with varying intensity (Fig. 7). In the case of fucoidan in solution (Fig. 7 A-D), frequent cytosolic calcium spiking was intensified by 50s after fucoidan addition. For platelets immobilized on fucoidan (Fig. 7 E-H) distinct cytosolic calcium spiking was observed. Comparison between fucoidan-induced calcium spiking between 25°C (Fig. 7 A) and 37°C (Fig. 7 C) shows that the frequency of spiking increases 3-fold at body temperature and there is also a 1.5-fold increase in amplitude. For platelets immobilized on VM64, at 37°C (Fig. 7 E) Ca^2+^ spiking frequency and intensity were increased substantially in comparison to 25 °C (Fig. 7G). This corresponds to the prediction that fucoidan-induced platelet activation could be influenced by the temperature dependent parameters: receptor translocation and enzyme turnover rates. A segregation of calcium spikes in groups appeared in all cells at 37 °C (Fig. 7C,G), but the origin of it is not yet clear. Further investigation to understand the basis of this behavior is needed. Together these data corroborate the model predictions that a) fucoidan induces calcium spiking in platelets and b) fucoidan-induced activation of platelets is significantly influenced by temperature. Spectral analysis of the spiking in spread on VM-64 antibodies demonstrated, that while height of the peaks corresponded at different temperatures, interspike interval was significantly shorter at 37°C than at 25°C. See supplementary video for additional details.

**Figure 7.**
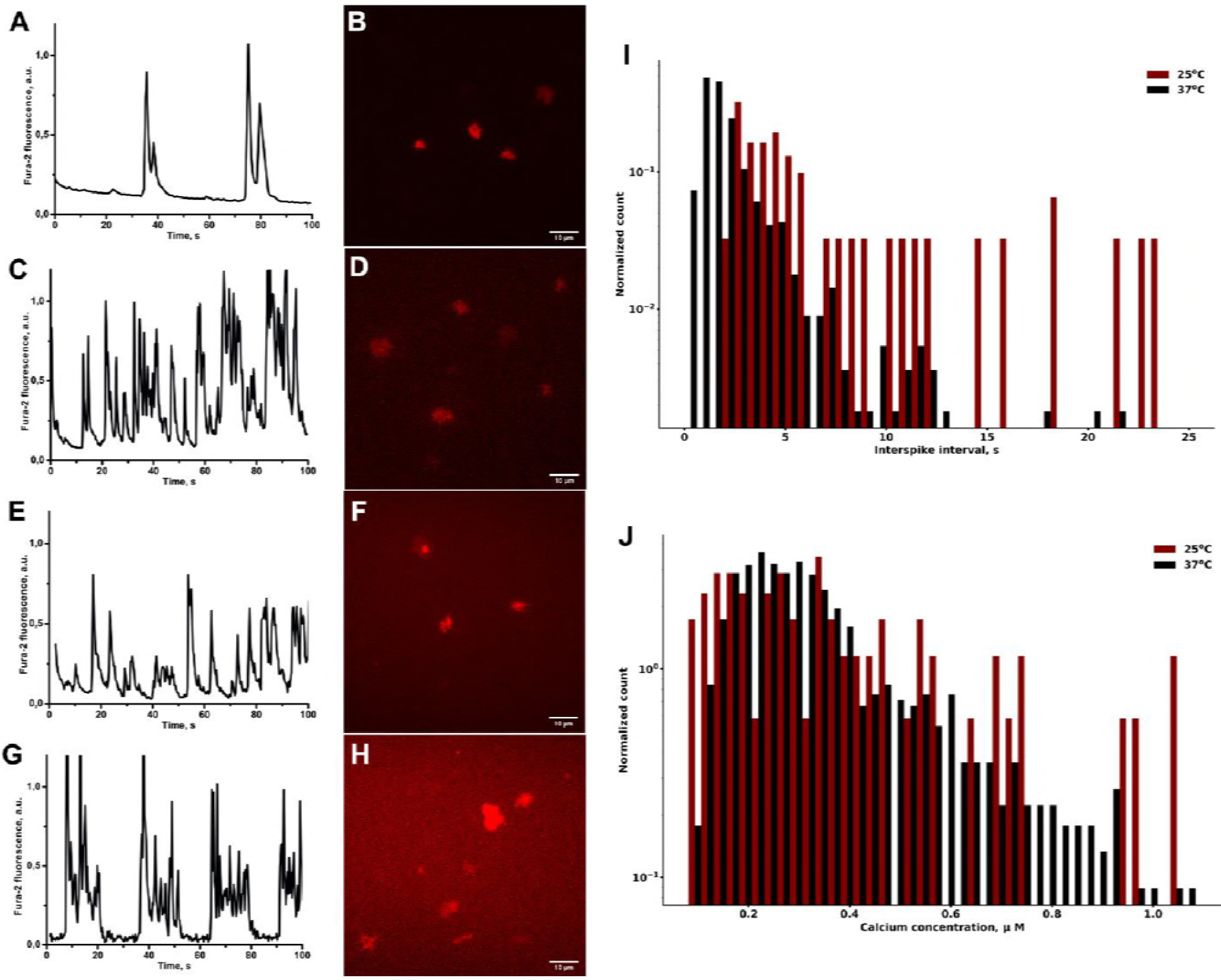
Cytosolic calcium spiking induced by fucoidan in single cells. Platelets were loaded with Fura-2, immobilized on VM-64 (A-D) or fucoidan (E-H) and then illuminated by 405 nm laser. Cytosolic calcium concentration was recalculated from Fura-2 fluorescence (see Methods). Cytosolic calcium spiking (A,C) and fluorescence (B,D) induced by fucoidan (200 μg/ml) immobilised on a cover glass. Thin layer fluorescence was monitored by TIRF microscopy at room temperature (A,B) or at 37 oC (C,D). Representative curves out of n=50. Washed platelets were immobilized on VM64 antibody (E-H). At time point “0” fucoidan solution at 100 μg/ml was added. Cytosolic calcium spiking (E,G) and whole cell fluorescence (F,H) were monitored at room temperature (E,F) or at 37 oC (G,H). Representative curves out of n=10. I,J – Intervals between calcium spikes (I) and calcium concentration per spike (J), spread on VM-64 and activated by fucoidan at 25°C (red) or at 37°C (black). Data collected from 20 cells.

## 3. Discussion

The objective of this study was to investigate the initial stages of CLEC-2 induced signalling in blood platelets. One of the main conclusions of our analysis is that, upon activation by various CLEC-2 ligands, CLEC-2 cytoplasmic domain phosphorylation is mediated by both SFK and Syk (Fig. 1, 3 A-C). We also conclude that a significant time delay observed between addition of CLEC-2 agonist and platelet activation may be explained by the rate-limiting clustering of CLEC-2 molecules in lipid rafts present within the platelet plasma membrane (Fig. 5). Despite this, activation of platelet CLEC-2 by fucoidan leads to cytoplasmic Ca^2+^ concentration oscillations (Fig. 6, 7).

Key to our investigation is the development of the first computational model of CLEC-2 induced platelet activation. The model describes the spatial distribution of CLEC-2 and its complexes with signalling enzymes in the plasma membrane after CLEC-2 ligation. *In silico* the formation of the LAT-signalosome within the lipid raft region appeared to be a crucial step in CLEC-2 signalling. Consequently, the rate of receptor translocation into the signalling region and the stability of lipid rafts were found to be rate-limiting factors in CLEC-2 induced platelet activation, alongside the activity of tyrosine kinases Syk, SFK and Btk (Fig. 3). Experimental measurements of cytosolic calcium concentration after platelet activation by CLEC-2 agonist fucoidan at different temperatures, and in the presence of a cholesterol sequestering agent mβCD, are in line with this conclusion.

Our study expands understanding of the significance of SFKs for CLEC-2 induced signalling in blood platelets. Previously, the role SFKs was often defined as the initial Syk activator (Hughes *et al*, 2015). An abrogation of CLEC-2 signalling following depletion of major mediators of SFK activation (PTP1B and CD148 (Mori *et al*, 2012)) as well as an inhibition of the CLEC-2 induced platelet activation by the broad SFK inhibitor PP2 were not yet explained. Based on the validation of different computational models we proposed that SFKs may phosphorylate CLEC-2 hemITAM in addition to Syk. This conclusion combines representations of Manne et al (Manne *et al*, 2015a) and Hughes et al (Hughes *et al*, 2015). Thus, the initial events in CLEC-2 signalling are as follows: First, bound to its ligand CLEC-2 is translocated to the signalling region (presumably, a lipid raft); Second, pre-phosphorylated by SFK on Y346, Syk kinase, as well as active SFK, phosphorylates CLEC-2 hemITAM; Third, both Syk and SFK bind to phospho-hemITAM in CLEC-2 by their SH2 domains; Fourth, Syk bound to CLEC-2 trans-autophosphorylates itself, or is phosphorylated by SFK and becomes catalytically active; Fifth, Syk phosphorylates LAT that leads to PLC2 incorporation and IP_3_ production (Fig. 1). The crucial role of Syk and SFK for these events was further confirmed by inhibitory analysis performed here (Fig. 3 A-C).

One of the aims of this work was to understand which mechanisms contribute to CLEC-2-induced platelet activation occurring much later than activation through GPVI, given that the intracellular signalling machinery of these two pathways are so similar (Dunster *et al*, 2015; Gibbins *et al*, 1997). The sensitivity analysis revealed that the initial steps in the CLEC-2-associated cascade are rate limiting (Fig. 3). To investigate the question further, we modified our computational model of CLEC-2 signalling cascade to describe the GPVI signalling cascade (Fig. S1). The rate-limiting steps of the GPVI model were tyrosine kinases Syk, SFK and Btk, which are characterized by reaction times of 10 s (Fig. S1 C-E), are significantly faster that the rate-limiting steps of the CLEC-2 cascade (lag-time 100 s, Fig. 3). Thus, we conclude that the differences in the initial steps in the CLEC-2 and GPVI cascades lead to CLEC-2 platelet activation being slower than activation through GPVI.

Our work leads to the proposition that temperature variation has a major effect on platelet activation through its action on the rate of receptor translocation into the signalling region, while its influence on enzymatic reactions is limited to catalytic constants. It is well known that the temperature dependence of turnover numbers of enzymes (Robinson, 2015; Struvay & Feller, 2012) is comparable with temperature dependence of membrane diffusion rates (Medda *et al*, 2015; Saha *et al*, 2015). Both CLEC-2 translocation rate and tyrosine kinase catalytic constants increase with temperature. Thus, the decrease in activation time in response to variation in temperature that we observed experimentally (Fig. 5) confirms the hypothesis that receptor molecule translocation in the membrane limits CLEC-2-induced platelet activation. Temperature variation had no major effect on ADP induced signalling, confirming the hypothesis that plasma membrane fluidity is specifically important for tyrosine-kinase signalling.

Our modelling predicts that the size of plasma membrane signalosome region has a large impact on CLEC-2 induced signal transduction (Fig. 3). This was confirmed experimentally here, when the depletion of cholesterol with mβCD, which makes any membrane microdomains less stable (Locke *et al*, 2002), led to significant increase in platelet activation lag-time (Fig. 5) in agreement with the previous work of Pollitt et al (Pollitt *et al*, 2010). This was modelled by the reduction of the size of lipid raft in the spatial model (Fig. S6, S7), which lead to a lack of CLEC-2 oligomerization and thus signalling abrogation, whereas the activation time increased with the decrease in raft radius. Yet, this hypothesis is not supported by the results of Manne et al (Manne *et al*, 2015b), where mβCD influenced only secondary platelet activation. However, in that work (Manne *et al*, 2015b) platelets were incubated with mβCD for an hour and this could have led to reintroduction of cholesterol in the signalling region, while dose-dependent response to mβCD, obtained upon incubation for 15 minutes in our work (Fig. S8 B) proves lipid rafts to be of high significance for prime CLEC-2 response.

Although our model predictions were supported by the experimental data, some limitations should be noted. First, the phosphatases in the model are not present explicitly (only CD148). For the stochastic model, no proper lipid raft discretization could be performed, thus, variation of the lipid raft size could not be assessed explicitly. The confirmations of model predictions here are limited to experiments with isolated platelets. A large part of the model predictions is concerned with kinase activity, which could not be assessed by single cell experiments. Furthermore, for inhibitory analysis of platelet tyrosine kinase signalling we performed experiments in the whole blood. Thus, distinction between primary and secondary signaling for tyrosine phosphorylation assays is complicated. Additional experiments on the roles of tyrosine kinases in CLEC-2 signalling as well as further model development (direct inclusion of the receptor clustering process, PTP1B phosphatase, lipid raft discretization) and investigation of cooperativity between CLEC-2 and GPVI should be the subject of further studies.

The fact that CLEC-2 activation is driven by the motion of proteins in the plasma membrane and the assembly of signalling complexes unveiled in this work, allows us to take a new perspective on all receptors that perform clustering after activation or are associated with specific lipid micro-domains. The knowledge of the underlying mechanisms of receptor cluster assembly will push forward understanding of molecular signalling in all types of eukaryotic cell.

## 4. Materials and methods

### 4.1. Reagents

The sources of the materials were as follows: calcium-sensitive cell-permeable fluorescent dye Fura-2-AM, Fura Red-AM, Fluo-3-AM and Fluo-4-AM (Molecular Probes, Eugene, OR); Fucoidan from *Fucus vesiculosis*, ADP, PGI_2_, EGTA, HEPES, bovine serum albumin, apyrase grade VII, methyl-β-cyclodextrin (mβCD) (Sigma-Aldrich, St Louis, MO); PRT060318 (MedChemExpress USA, Monmouth Junction, NJ); PP2 and PP3 (Tocris Bioscience; Ellisville, MO, USA). VM-64 antibody was a kind gift of Dr. A.V. Mazurov (NMRC of Cardiology, Moscow, RF) (Mazurov *et al*, 1991). Rhodocytin was a kind gift of Dr. A. Pollitt (ICMR, Reading University, Reading, UK) (Severin *et al*, 2011).

### 4.2. Blood collection and platelet isolation

Healthy volunteers, both men and women aged between 18 and 35 years were recruited into the study. Investigations were performed in accordance with the Declaration of Helsinki, and written informed consent was obtained from all donors. Blood was collected into 4.5 ml tubes containing 3,8% sodium citrate (1:9 vol/vol) and supplemented by apyrase (0.1 U/mL). Platelets were purified by double centrifugation as described previously (Panteleev *et al*, 2005; Sveshnikova *et al*, 2016). Briefly, platelet-rich plasma was obtained by centrifugation at 100 g for 8 minutes. Platelet-rich plasma was supplemented with additional sodium citrate (27 mM) and centrifuged at 400 g for 5 minutes. The resultant supernatant was removed and platelets resuspended in Tyrode’s buffer (150 mM NaCl, 2.7 mM KCl, 1 mM MgCl2, 0.4 mM NaH2PO4, 5 mM HEPES, 5 mM glucose, 0.2% bovine serum albumin, pH 7.4). Alternatively, blood was collected in Li-heparine (IMPROVACUTER^®^) or hirudin (SARSTEDT Monovette^®^) containing vacuum tubes.

### 4.3. Flow cytometry and inhibitory analysis

For continuous flow cytometry experiments, washed platelets were incubated with either 2 μM Fura Red-AM (or 2 μM of Fluo-3 or Fluo-4 and 2 μM of Fura-2) prior to the final wash for 45 minutes at room temperature or for 30 min at 37°C in the presence of apyrase (1 U/mL). Platelets were then incubated in buffer A for 10 minutes and then centrifuged. Whole blood was incubated with either 2 μM Fura Red-AM (or 2 μM of Fluo-3 or Fluo-4 and 2 μM of Fura-2) for 30 min at 37°C in the presence of apyrase (1 U/mL). Whole blood was diluted 20-times with calcium and albumin containing Tyrode’s buffer. Samples were diluted to concentration 1000 plt/μl and analyzed using FACS Canto II or FACS Aria (BD Biosciences, San Jose, CA, USA) flow cytometer in a continuous regime with 20s interruption for the addition of an activator. For inhibitory analysis hirudinated platelet rich plasma was incubated with either 20 μM PP2 or PP3 (inactive analogue control) or 5 μM PRT060318.

### 4.4. Immunoblotting

Human platelets from drug-free volunteers were prepared on the day of the experiment as described previously (Gibbins, 2004) and suspended in modified Tyrodes-Hepes buffer (134 mM NaCl, 0.34 mM Na_2_HPO_4_, 2.9 mM KCl, 12 mM NaHCO_3_, 20 mM Hepes, 5 mM glucose, 1 mM MgCl_2_, pH 7.3) to a density of

1.5 × 10^9^ cells/ml. Stimulation of platelets with 10x fucoidan (final concentration: 1 μg/ml 10 μg/ml, 100 μg/ml), was performed for 0-15-30-60-90-120-150-180-300 s at 25 or 37°C in an aggregometer with continuous stirring (1000 rpm). Reactions were abrogated by addition of 4x SDS-PAGE sample treatment buffer (200 mM Tris-HCl pH6.8, b-MeEtOH 400 mM, SDS 4%, Bromphenol blue 0.01%, Glycerol 40%). Samples were then heated to 99°C for 10 minutes and centrifuged at 15000g for 10 minutes in order to remove cell debris.

Proteins were separated by SDS-PAGE on 10% gels and transferred to polyvinylidene difluoride (PVDF) membranes that were then blocked by incubation in 5% (w/v) bovine serum albumin dissolved in TBS-T. Primary and secondary antibodies were diluted in TBS-T containing 2% (w/v) bovine serum albumin and incubated with PVDF membranes for 1.5 h at room temperature. Blots were washed 4 times for 15 minutes in TBS-T after each incubation with antibodies and then developed using an enhanced chemiluminescence detection system using ECL Prime western blotting detection reagent. Primary antibodies were used at a concentration of 1 μg/ml (anti-phosphotyrosine PY20) or diluted 1:1000 (anti-tubulin). Horseradish peroxidase-conjugated secondary antibodies were diluted 1:1000. In order to control for protein loading, membranes were stripped by washing 2 times for 30 minutes in stripping buffer (250 mM Glycine, 0.2% SDS, 0.1% Tween-20, pH 2.2) twice for 10 minutes in PBS and 2 times for 5 minutes in TBST at room temperature. Membranes were then blocked for 30 minutes by 2% TBS-T BSA solution at room temperature and re-stained with anti-tubulin antibodies.

### 4.5. Depletion of platelet cholesterol

Cholesterol was depleted from the plasma membrane of platelets, by incubation of washed platelets for 15 minutes with different concentrations of mβCD at 37° C before stimulation as described in (Mahammad & Parmryd, 2015).

### 4.6. Microscopy

For microscopy experiments, platelets were loaded with calcium fluorophores and immobilized by either incubation of platelet suspension in flow chamber for 5 min, or by perfusing whole blood over the surface at a shear rate of 200 s^−1^ for 5 minutes. For total internal reflection fluorescent (TIRF) microscopy, platelets were immobilized either on fucoidan (100 μg/ml) or anti-CD31 (VM-64) (Mazurov *et al*, 1991) and investigated in flow chambers (Lawrence *et al*, 1987). An inverted Nikon Eclipse Ti-E microscope equipped with 100x/1.49 NA TIRF oil objective was used. Cells were observed in DIC and TIRF modes. 405 nm laser was applied to assess calcium-free Fura-2 fluorescence alternatively with 488 nm laser for calcium-bound Fluo-3 fluorescence in a platelet. Calcium concentration was assessed either from the ratio of fluorescence of Fluo-3 and Fura-2 according to one-Kd formula (Sveshnikova *et al*, 2016) or from Fura-2 fluorescence as a ratio of initial and running values with taking exponential bleaching of the dye into account. For temperature fixation during observation a lens heater (Bioptechs, Butler, PA) was used.

### 4.7. Data analysis

Nikon NIS-Elements software was used for microscope image acquisition; ImageJ (http://imagej.net/ImageJ) was used for image processing for both TIRF-microscopy and western blotting assays from literature. Flow cytometry data was processed using FlowJo (http://www.flowjo.com/) software. Statistical analysis was performed in Python 3.0.

### 4.8. Model solution

The reaction-diffusion model, formed of partial differential equations (see supplement) with initial variable values (Table S3, S4, S6, S7) was integrated using the method of Lines and CVODE in VCell software (www.VCell.org, access: **AlleMart:** CLEC-2 v5.1 Syk-Src, **AlleMart:** CLEC-2 v5.1 Syk). The stochastic model was solved using stochastic integration methods (the tau-leap method (Pahle, 2009; Gillespie, 2007)) implemented in COPASI software, in a similar manner to previously published methods (Balabin & Sveshnikova, 2016; Sveshnikova *et al*, 2016). Stochastic models can be found in supplementary (See Table S9 supplementary files).

## Acknowledgements

We thank Prof. F.I. Ataullakhanov (CTP PCP RAS, Moscow, RF), Dr. A.V. Mazurov (NMRC of Cardiology, Moscow, RF) and Dr. N.E Ustuzhanina (ZIOC RAS, Moscow, RF) for reagents used during preliminary experiments and valuable discussions. We are grateful to Dr. A.V. Pichugin (FMBA, Moscow, RF) for advice and Miss V.N. Kaneva for assistance during flow cytometry data collection.

## 5. Authors contributions

A.A.M. developed the model, performed simulations, performed experiments (flow cytometry, immunoblotting), analyzed the data and wrote the paper. F.A.B. performed single-cell microscopy experiments. J.M.G. and J.L.D. analyzed the data and edited the paper. M.A.P. supervised the project and edited the paper. A.N.S. planned model development and research, analyzed the data, performed experiments (microscopy) and edited the paper. The authors declare that they have no conflict of interest.

## Abbreviations

CLEC-2: C-type lectin-like receptor 2
LEC: lymphatic endothelial cells
GPVI: glycoprotein VI
FcR: Fc receptor
ITAM: immune tyrosine activation motif
Syk: spleen tyrosine kinase
SFK: Src-family kinase
PI3K: phosphoinositide 3-kinase
PIP_2_: phosphatidylinositol 4,5-bisphosphate
PIP_3_: phosphatidylinositol 3,4,5-trisphosphate
LAT: linker adapter for T-Cells
PLCɣ2: phospholipase Cɣ
PGI_2_: prostaglandin I_2_
mβCD: methyl β cyclodextrin
TIRF: total internal reflection
PTP1B: protein tyrosine phosphatase 1B
Btk: Bruton’s tyrosine kinase
TxA2: Thromboxane A2

